# Genetic analysis of isoform usage in the human anti-viral response reveals influenza-specific regulation of *ERAP2* transcripts under balancing selection

**DOI:** 10.1101/188961

**Authors:** Chun Jimmie Ye, Jenny Chen, Alexandra-Chloé Villani, Rachel E. Gate, Meena Subramaniam, Tushar Bhangale, Mark N. Lee, Towfique Raj, Raktima Raychowdhury, Weibo Li, Noga Rogel, Sean Simmons, Selina H. Imboywa, Portia I. Chipendo, Cristin McCabe, Michelle H. Lee, Irene Y. Frohlich, Barbara E. Stranger, Philip L. De Jager, Aviv Regev, Tim Behrens, Nir Hacohen

## Abstract

While the impact of common genetic variants on gene expression response to cellular stimuli has been analyzed in depth, less is known about how stimulation modulates the genetic control of isoform usage. Analyzing RNA-seq profiles of monocyte-derived dendritic cells from 243 individuals, we uncovered thousands of unannotated isoforms synthesized in response to viral infection and stimulation with type I interferon. We identified more than a thousand single nucleotide polymorphisms associated with isoform usage (isoQTLs), > 40% of which are independent of expression QTLs for the same gene. Compared to eQTLs, isoQTLs are enriched for splice sites and untranslated regions, and depleted of sequences upstream of annotated transcription start sites. Both eQTLs and isoQTLs in stimulated cells explain a significant proportion of the disease heritability attributed to common genetic variants. At the *IRF7* locus, we found alternative promoter usage in response to influenza as a possible mechanism by which DNA variants previously associated with immune-related disorders mediate disease risk. At the *ERAP2* locus, we shed light on the function of the major haplotype that has been maintained under long-term balancing selection. At baseline and following type 1 interferon stimulation, the major haplotype is associated with absence of *ERAP2* expression while the minor haplotype, known to increase Crohn’s disease risk, is associated with high *ERAP2* expression. Surprisingly, in response to influenza infection, the major haplotype results in the expression of two uncharacterized, alternatively transcribed, spliced and translated short isoforms. Thus, genetic variants at a single locus could modulate independent gene regulatory processes in the innate immune response, and in the case of *ERAP2*, may confer a historical fitness advantage in response to virus.

## Introduction

An important aspect of eukaryotic gene regulation is the usage of alternative gene isoforms. This is achieved through several mechanisms at the transcript level, including: alternative promoters for transcription initiation, alternative splicing of pre-messenger RNA, alternative polyadenylation, and selective degradation of isoforms. These processes regulate the relative abundances of multiple coding and non-coding RNAs from the same underlying DNA sequence, often resulting in altered function of the gene products in response to developmental or environmental changes (Black 2003; Maquat 2004; Matlin et al. 2005; Wang et al. 2008).

A case in point is the role of alternative isoform usage in the human immune response. For example, studies have shown that alternative splicing is critical across many immune processes, such as the balance between IgM and IgD immunoglobulin isoforms in B cells (Enders et al. 2014), naïve and memory T cell proportions controlled by *CD45* isoforms (Berard and Tough 2002), and innate immune responses to pathogens regulated by different isoforms of *MYD88* (Martinez and Lynch 2013). Genetic variants that affect isoform usage have been associated with immune disorders (Xiong et al. 2015) including single nucleotide polymorphisms (SNPs) that alter the relative splicing of two *IRF5* isoforms (Graham et al. 2007) increasing the risk of systemic lupus erythematosus (SLE).

Previous studies have identified shared and divergent transcriptional programs in the antibacterial and antiviral responses of innate immune cells (Amit et al. 2009; Lee et al. 2014), with genetic variants imparting stimulation specific effects on the total transcript abundances of thousands of genes (Barreiro et al. 2012; Fairfax and Knight 2014; Lee et al. 2014; Quach et al. 2016). While maps of genetic determinants of alternative isoform usage are beginning to emerge, most notably in lymphoblastoid cell lines (Lappalainen et al. 2013; Li et al. 2016), across post-mortem human tissues (Consortium 2015; Rivas et al. 2015), and in macrophages stimulated with bacteria (Nedelec et al. 2016), differential isoform usage in the human antiviral response, its natural variability, and its genetic basis have not been studied.

Here, we integrate bulk RNA-sequencing with dense genotyping to systematically investigate the genetic control of isoform usage in monocyte derived dendritic cells (MoDCs) at rest, and in response to influenza-infection or type 1 interferon. Because the type 1 interferon pathway is known to be engaged by a broad array of microbial products, our study design is unique in allowing the separation of the universal and influenza-specific interferon-induced responses. Since the human transcriptome has never been annotated under these conditions, we first used *de novo* assembly to catalog and expectation maximization to quantify all synthesized isoforms in resting and stimulated MoDCs. Then, by harnessing the natural transcriptomic and genetic variation in the ImmVar cohort (Lee et al. 2014; Raj et al. 2014a; Ye et al. 2014), we mapped genetic variants (isoQTLs) associated with alternate isoform usage. Systematic characterization of isoQTLs, especially in comparison to eQTLs, provides mechanistic insights into the genetic control of different aspects of gene regulation and enables the functional interpretation of loci associated with immune-related diseases and under natural selection.

## Results

### Influenza and type I interferon stimulate widespread alternate isoform usage

We used paired-end RNA-seq to profile the transcriptomes of primary MoDCs from healthy donors at rest (N = 99), and following stimulation with either influenza ΔNS1 (a strain engineered to maximize the IFNβ-induced host response to infection by the deletion of a key virulence factor (Shapira et al. 2009)) (N = 250) or interferon beta (IFNβ), a cytokine that stimulates anti-viral effectors (N = 227). A total of 552 pass-filter samples (out of 576), 84 from all three conditions, 127 from both stimulation conditions, and 46 from only one condition, were analyzed (**Table S1**). To define the corpus of transcribed isoforms in human MoDCs at rest and in response to stimulation, including previously unannotated isoforms, we assembled the transcriptome *de novo* in each sample (individual-condition pair) from RNA-seq alignments, retained only expressed isoforms (> 5 transcripts per million (TPM) in any sample), then combined isoforms across all samples to enable direct comparisons between conditions. We note that unannotated isoforms that do not match current Gencode, UCSC, or RefSeq annotations (Harrow et al. 2006; O’Leary et al. 2016; Casper et al. 2018) were enriched in genes expressed at less than 25 TPMs across all three conditions (**Fig. S1**) and thus removed these genes and their corresponding transcripts from downstream analyses.

Our final assembled transcriptome contains 15,754 transcripts corresponding to 8,194 genes (**Table S2**), 64.5% of which have transcriptional structures that exactly match an annotated transcript. 5,204 transcripts (33.0%) contain a novel splice site and 389 transcripts (2.5%) do not harbor novel splice sites but contain novel splice junctions. These novel splice sites and splice junctions are well supported by the presence of spliced reads (> 10 mapped reads) spanning exon junctions of our assembled transcriptome. Of the 25,099 novel splice events, 11,093 (44%) are within 5’ UTRs and 7,251 (28.9%) are within 3’ UTRs, echoing previous deep sequencing analyses reporting the majority of novel isoforms are due to alternative splicing of UTRs (Deveson et al. 2018). An additional 2,663 transcripts (16.9%) are annotated with a novel transcription start site (TSS). To assess the accuracy of these new TSSs, we aligned CAGE sequencing reads from resting MoDCs (Noguchi et al. 2017) to the set of unique TSSs (TSS + 500 bp downstream) from assembled transcripts in the baseline condition. Compared to all TSSs, 69% (vs 84%) of the new TSSs were supported by the presence of at least 1 mapped CAGE read and 40% (vs 61%) were supported by > 5 mapped reads. By visual inspection, we find the most common mis-annotation to be isoform reconstructions that begin within the gene body, downstream of the annotated TSS. While some of these fragments may truly exist, we are unable to verify them with short read sequencing and thus conservatively estimate the false annotation rate to be ~20% (difference between new and all TSSs) of new TSSs or 3.3% of our entire transcriptome.

We next compared changes in isoform usage – estimated as the ratio of isoform abundance over total gene abundance - in response to each stimulus. Relative to baseline, the usage of ~2x as many isoforms (5,326 vs 2,509) were altered in flu-infected compared to IFNβ-stimulated cells (beta regression, FDR < 0.05; **Table S3, Fig. 1A**). This is corroborated by directly comparing flu-infected and IFNβ-stimulated cells where the usage of 5,072 isoforms were altered (beta regression, FDR < 0.05; **Table S3, Fig. S2**). In response to either stimuli, more than a third of the isoforms with differential usage were previously unannotated, highlighting the inadequacy of current transcriptome annotations in describing the full repertoire of gene isoforms in the human antiviral response. Of the differentially expressed genes (DESeq, FDR < 0.05, gene abs(log_2_[fold change]) > 1, **Table S4**) with more than one isoform, 54% (2,233/4,120) in flu-infected cells and 29% (1,122/3,898) in IFNβ-stimulated cells had at least one isoform that differed in usage (**Fig. 1A**).

**Figure 1.**
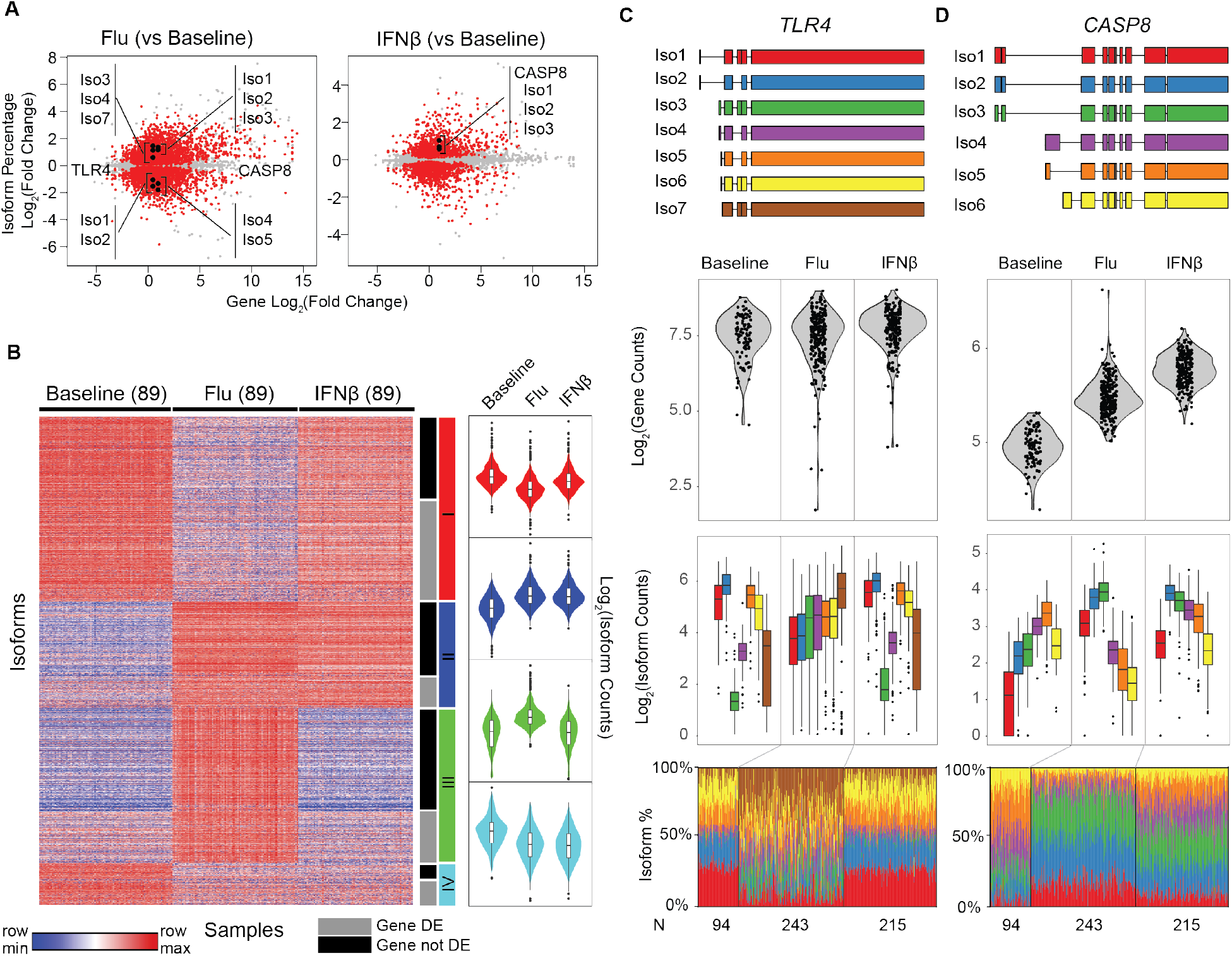
Transcriptome changes in response to stimulation. (**A**) Scatter plot of log_2_ fold change of gene abundance (x-axis) versus log_2_ fold change of isoform usage (y-axis) in flu-infected (left) and IFNβ-simulated (right) cells compared to baseline. Each dot represents one isoform. Isoforms that significantly differed in their usage (beta regression, FDR < 0.05) are highlighted in red. (**B**) Clustering of isoform usage percentages in baseline, flu-infected, and IFNβ-stimulated cells. Heatmap colors are row scaled (red: row max, blue: row min) (left). Violin plots (right) summarize the usages of all isoforms within a cluster separated by condition. Only isoforms (one per gene) that most significantly changed (beta regression, FDR < 0.05) in usage are shown. (**C** and **D**) *De novo* constructed isoforms (top panel), gene abundance (middle panel), and isoform abundance and usage (bottom panels) for (**C**) *TLR4* and (**D**) *CASP8*.

Isoforms that differed in usage partitioned into four prominent clusters (**Fig. 1B**, k-means clustering of the most significant isoforms, one per gene, **Table S5**). Isoforms with increased usage in response to both stimuli (Cluster II) were highly enriched for innate system processes (GO:0002376, q < 6.41 ×10^−6^), defense response to virus (GO:0051607, q < 9.56×10^−6^), and type 1 interferon signaling pathway (GO:0060337, q < 1.25 × 10^−3^). Isoforms with increased usage in response to flu but not IFNβ (Cluster III) were enriched for regulators of gene expression (GO:0010468, q < 6.23× 10^−3^) including genes involved in the regulation of MAP kinase cascade (GO:0043408, q < 1.13×10^−2^) and inflammatory response (GO:0006954, q < 1.88×10^−2^). Isoforms with decreased usage in response to flu (Cluster I) were enriched for oxidoreductase activity (GO:0016616, q < 2.47×10^−2^); isoforms with decreased usage in response to both conditions (Cluster IV) were not significantly enriched for known gene ontology entries. *TLR4*, the toll-like receptor classically associated with sensing bacterial ligands but have been shown to also sense viral products (Doyle et al. 2002), is among the genes that have flu-specific isoform usage despite little change in total gene abundance (**Fig. 1C, Fig. S3**). In flu-infected cells, the usages of longer isoforms with an upstream alternative start site (*TLR4*/Iso1 and *TLR4*/Iso2) were decreased, while the usages of *TLR4*/Iso3, *TLR4*/Iso4, and *TLR4*/Iso7 were increased. *TLR4*/Iso4 encodes the annotated 839 aa product, while isoforms *TLR4*/Iso3 and *TLR4*/Iso7 encode shorter, 799 aa products each with a truncated extracellular domain missing a predicted signal peptide. We also found decreased usage of short *CASP8* isoforms (*CASP8*/Iso4, *CASP8*/Iso5, *CASP8*/Iso6, **Fig. 1D, Fig. S3**) only in flu-infected cells. Although *CASP8* is known to induce the apoptotic program via the Fas-associated death domain (FADD) protein in response to extrinsic cytokine signals, *CASP8*/Iso4 has a unique N-terminal extension of 59 amino acids, which has been reported to allow for selective recruitment to the endoplasmic reticulum (Breckenridge et al. 2002). These results demonstrate that changes in isoform usage independent of overall gene abundance are pervasive and affect prominent innate immune sensors and regulators in viral versus interferon response.

### Genetic variants associated with isoform usage are enriched for distinct gene regulatory elements

While it is known that common genetic variants modulate gene expression (Lee et al. 2014), we assessed if they could also affect isoform usage in both resting and stimulated MoDCs. We associated over 10 million (M) imputed variants with two transcriptional traits, isoform percentage and total gene abundance, to identify isoform usage quantitative trait loci (isoQTLs) and expression quantitative trait loci (eQTLs), respectively. After adjusting for unwanted variation that likely track with technical and biological confounders (**Fig. S4** and **S5**), we identified 2,763 isoforms corresponding to 1,425 genes (linear regression, permutation FDR < 0.05, **Table S6**) with local isoQTLs (+/- 500kb of TSS) and 6,694 genes (linear regression, permutation FDR < 0.05, **Table S7**) with local eQTLs in at least one condition. A substantial proportion of leading isoQTL SNPs (63% baseline, 40% flu, 41% IFNβ) were not significant eQTLs, suggesting that the genetic control of isoform usage and overall gene abundance are largely independent.

Genetic variants could modulate isoform usage through several mechanisms including perturbing the usage of alternate promoters, splice sites, or regulatory elements in the UTRs. We compared the genomic properties of isoQTLs and eQTLs to identify the mechanisms by which each class of variants acts. When normalized by exon and intron lengths, leading SNPs for local isoQTLs (one per isoform) were enriched across the entire gene body (**Fig. 2A**), in distinct contrast to leading SNPs for local eQTLs (one per gene), which were enriched near TSSs and transcription end sites (TESs). Further, when compared to a set of eQTLs matched for allele frequency and distance to TSS, leading SNPs for local isoQTLs were most enriched for splice sites (baseline: 3.7x, flu: 3.0x, IFNβ: 4.0x), synonymous (baseline: 1.9x, IFNβ: 2.1x) and missense variants (baseline: 1.9x, flu: 1.8x), and 5’ UTRs (baseline: 1.3x, IFNβ: 1.2x) (**Fig. 2B**). Compared to eQTLs, isoQTLs are not enriched for binding sites of transcription factors known to play a role in myeloid cell response (**Fig. S6**). These results suggest that genetic variants associated with isoform usage likely do so via *cis* regulatory sequences that modulate alternative splicing and transcript stability.

**Figure 2.**
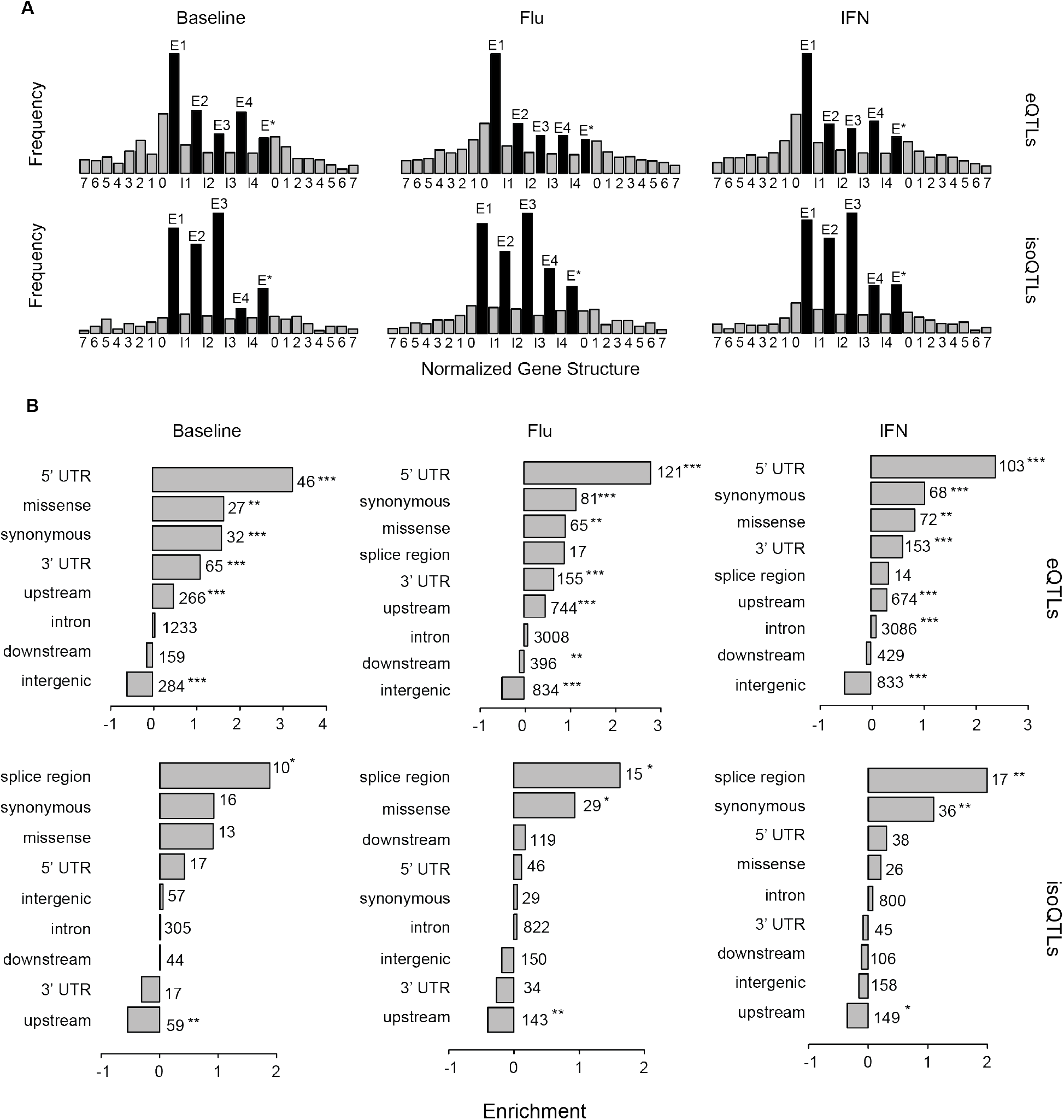
Properties of local eQTLs and isoQTLs. (**A**) Frequency (y-axis) of the location of leading SNPs for local eQTLs and isoQTLs (permutation FDR < 0.05) with respect to meta gene structure (x-axis). Genes are normalized to five exonic regions (E1 - E4: exons 1 - 4, E*: exon 5 - last exon) and four intronic regions (introns 1, 2, 3, and 4 – last intron). Upstream and downstream sequences are divided into 100kb windows. (**B**) Log_2_ fold enrichment (x-axis) of leading SNPs for local eQTLs and isoQTLs (permutation FDR < 0.05) for genomic annotations. EQTL enrichments are calculated using a background set of SNPs matched for distance to TSS and allele frequency. IsoQTL enrichments are calculated with respect to a background set of eQTLs matched for distance to TSS and allele frequency. Multiple testing significance is indicated (*: P < 0.05, **: P < 0.01, ***: P < 0.001).

### Genetic control of alternative isoform usage in responses to virus and interferon

To assess how the genetic control of isoform usage differs in response to stimuli, we analyzed 84 donors whose cells were assayed in all three conditions to enable equally powered comparisons across conditions. At these sample sizes, we detected more eQTLs in cells stimulated with IFNβ than cells at rest or infected with flu (baseline: 1,715, flu: 1,755, IFNβ: 2,108, permutation FDR < 0.05, **Table S6**) but similar numbers of isoQTLs across conditions (baseline: 717, flu: 644, IFNβ: 692, permutation FDR < 0.05, **Table S7**). For the 1,164 isoforms with isoQTLs in at least one condition, we compared the effect sizes (R_iso_^2^) of associations across conditions (**Fig. 3A**). The correlation of R_iso_^2^s was lowest between flu-infected and resting (baseline) cells (Pearson ρ_flu.baseline_=0.61 compared to ρ_IFN.baseline_=0.76 and ρ_IFN.flu_=0.74). The corresponding genes of isoforms with higher R_iso_^2^ in stimulated cells were upregulated in response to stimuli, suggesting that the genetic control of isoform usage is sensitive to activation of gene regulatory programs that control overall gene abundance (**Fig. 3AB**).

**Figure 3.**
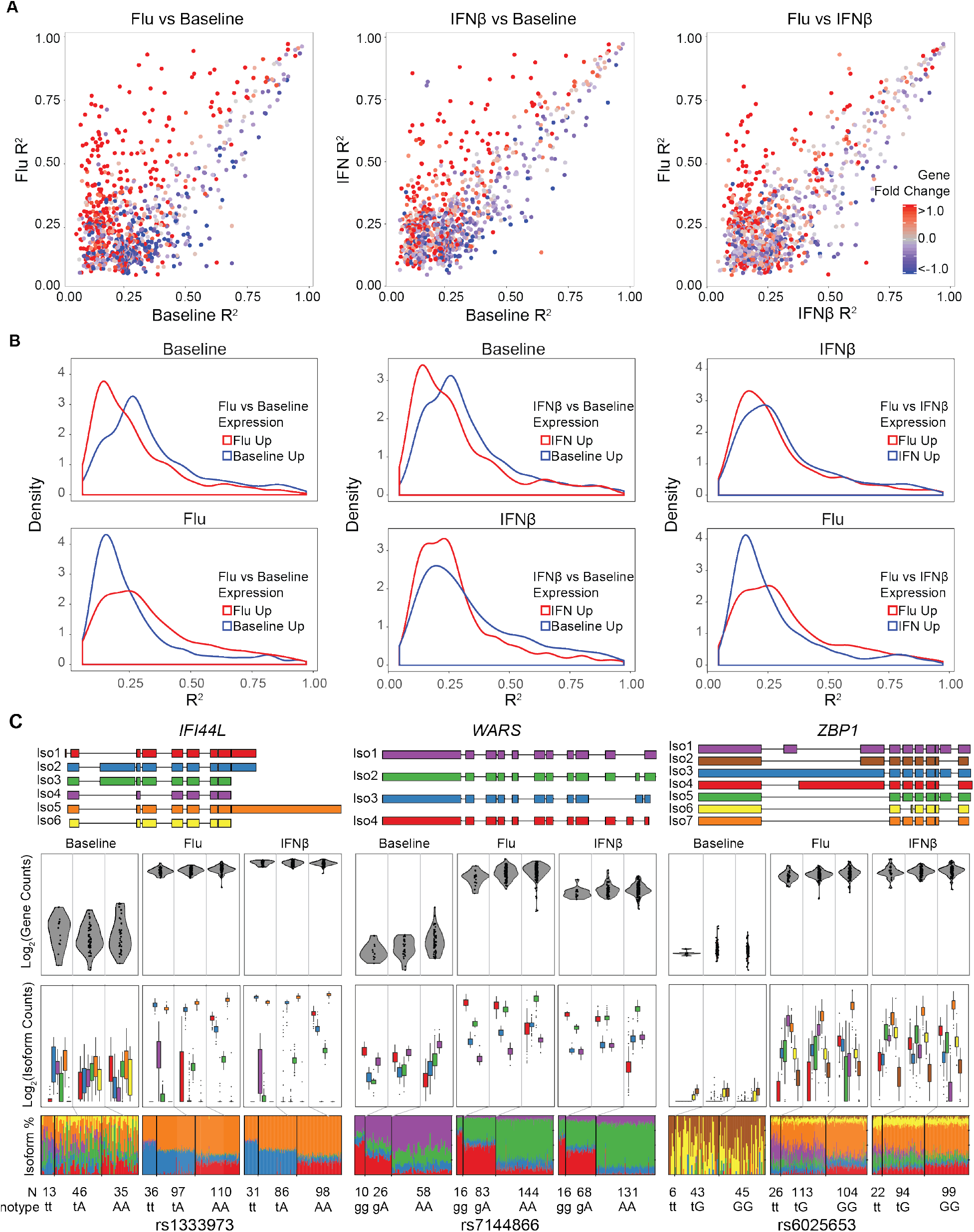
Comparison of local isoQTLs between conditions. (**A**) Correlation of effect sizes (R^2^) for significant local isoQTLs (permutation FDR < 0.05) between pairs of conditions. Transcripts are colored by differential expression (red: up-regulated in condition 2, y-axis, blue: up-regulated in condition 1, x-axis) for each pair of conditions. (**B**) Distributions of effect sizes (R^2^) for significant local isoQTLs (permutation FDR < 0.05) for each pair of conditions segregating genes based on expression in each condition. (**C**) *De novo* constructed transcript structure (top panel) and box-whisker plots (bottom 3 panels) between transcript quantitative traits (y-axis: log2(normalized gene abundance), normalized isoform abundance, or isoform percentage) and genotype (x-axis) for 3 genes (*IFI44L, WARS* and *ZBP1*) with risoQTLs.

To directly assess how stimulation modifies the effects of genetic variants on isoform usage, we mapped SNPs associated with the difference in isoform usage between conditions, herein referred to as local response-isoQTLs (risoQTLs). Compared to resting cells, we identified 53 (flu) and 30 (IFNβ) isoforms, corresponding to 31 and 14 genes, with at least one local risoQTLs (permutation FDR < 0.05, **Table S8**). Amongst the 7 genes that share local risoQTLs in both stimulated conditions were *IFI44L* and *WARS* (**Fig. 3C**), two genes whose splicing has previously been studied. *IFI44L* is a type 1 interferon-stimulated gene known to moderately inhibit human hepatitis virus replication *in vitro* (Schoggins et al. 2011) and whose splicing has been shown to be influenced by rs1333973 (Lalonde et al. 2011). Rs1333973 is the most significant risoQTL associated with the usages of two isoforms differentiated by exon 2 in flu-infected and IFNβ-stimulated cells (flu vs baseline: isoform 1: +8.9%, P < 1.37×10^−9^, isoform 2: −13.9%, P < 1.53×10^−9^; IFNβ vs baseline: isoform 1: +9.6%, P < 1.21×10^−10^, isoform 2: −15.7%, P < 2.14×10^−10^, **Fig. S7**). *WARS* is a tryptophanyl-tRNA synthetase primarily involved in protein synthesis. While *WARS* isoforms are known to encode for catalytic null enzymes (Lo et al. 2014) and have anti-angiogenic activity inducible by IFNγ (Wakasugi et al. 2002), there has been no previous reports of the genetic control of these isoforms. The most significant risoQTL rs7144866 is associated with the usages of two isoforms differentiated by exon 2: while rs7144866^A^ increases the usage of isoform 1 in cells at rest, it increases the usage of isoform 2 in flu-infected and IFNβ-stimulated cells resulting in the risoQTL (flu vs baseline: −19.1%, 2.50×10^−29^, ifn: −17.7%, 7.03×10^−26^, **Fig. S8**). There were 40 isoforms, corresponding to 30 genes, that have at least one risoQTL in flu-infected cells compared to interferon-stimulated cells (permutation FDR < 0.05, **Table S8**). Amongst these was *ZBP1*, a sensor of influenza infection that triggers cell death and inflammation and contributes to virus-induced lethality (Kuriakose et al. 2016). We found rs6025653^t^ increases the usage of isoform 1, which is differentiated from all other isoforms by the retention of exon 9, by 9.67% (P < 4.16× 10^−16^, **Fig. 3C, S9**) in flu-infected compared to interferon-stimulated cells. These results suggest that while influenza-infected and interferon-stimulated cells share some genetic control of isoform usage reflecting a common gene regulatory program (as interferons are induced by viral infection), influenza infection confers specific genetic control of isoform usage in previously unknown viral sensing pathways independent of downstream effector (type-1 interferon) signaling.

### Association of eQTLs and isoQTLs with immune-related diseases

Previous analyses of the overlap between expression QTLs and genome-wide association studies (GWAS) have aided the localization and functional interpretation of causal variants in GWAS loci. Because disease-causing variants that affect isoform usage could have more profound effects on gene regulation by altering protein structure, we jointly analyzed disease-associated variants and local isoQTLs in addition to local eQTLs using two approaches. First, compared to SNPs from the latest GWAS catalog (MacArthur et al. 2017), local eQTLs in stimulated cells were enriched in multiple diseases including inflammatory bowel disease (flu P < 5.46×10^−7^, IFNβ P < 5.21 ×10^−5^), rheumatoid arthritis (flu P < 2.49×10^−5^, IFNβ P < 0.03), and Parkinson’s disease (flu P < 4.06×10^−7^, IFNβ P < 7.44×10^−5^) (**Fig. S10**), while local isoQTLs were enriched in late onset Alzheimer’s disease (flu P < 1.15×10^−6^, IFNβ P < 1.75×10^−4^), vitiligo (flu P < 4.94×10^−5^, IFNβ P < 0.42), and SLE (flu P < 0.1, IFNβ P < 3.62×10^−3^) (**Fig. S10**). Notably, the overlap of isoQTLs with Alzheimer’s loci captures genes with known variants that affect splicing (*CD33* (Hernandez-Caselles et al. 2006; Raj et al. 2014b), *CD46* (Russell et al. 1992)). Corroborating this, we performed partitioned heritability analysis using linkage disequilibrium (LD) score regression (Finucane et al. 2015). For 28 traits with available summary statistics, both local eQTLs and local isoQTLs explained a statistically significant percentage of the SNP heritability of autoimmune diseases (i.e. ulcerative colitis, SLE), some neurodegenerative diseases (i.e. Alzheimer’s), and did not explain much of the SNP heritability of diseases with no known relationship to the innate immune system (i.e. Type 2 diabetes) (**Fig. 4A, Table S9**). These results suggest a role for both variants that affect isoform usage and gene expression in mediating autoimmune and neurodegenerative disease risk.

**Figure 4.**
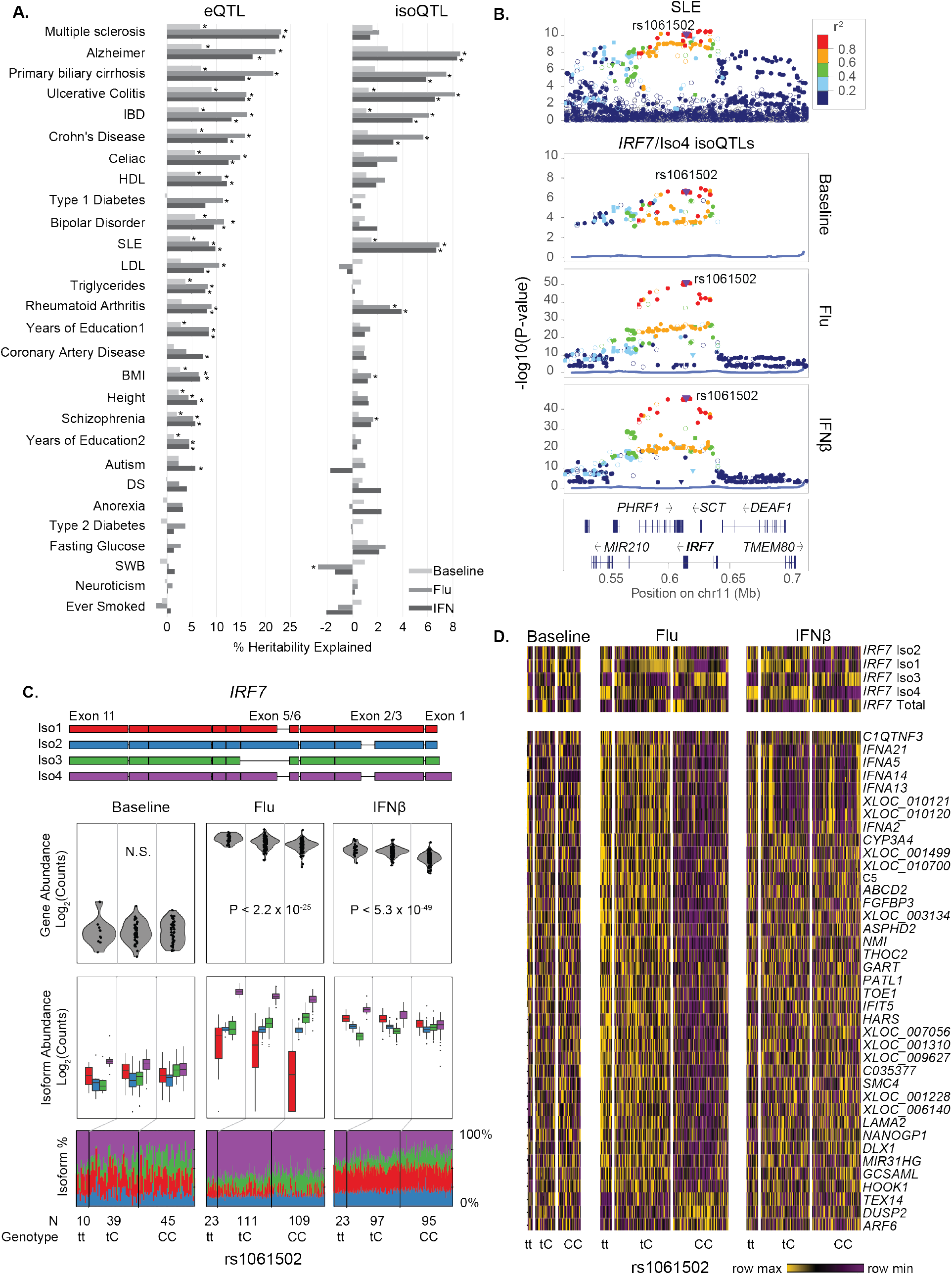
GWAS enrichment of local eQTLs and isoQTLs. (A) Partitioned heritability analysis showing the proportion of SNP heritability explained (x-axis) for 28 traits (y-axis) by eQTLs (left) and isoQTLs (right). DS = depressive symptoms, SWB = subject well being, IBD: inflammatory bowel disease, SLE: systemic lupus erythematosus, BMI: body mass index, HDL: high density lipoprotein, LDL: low density lipoprotein. (B) LocusZoom plots of the *IRF7* region for SLE GWAS associations (top panel) and *IRF7*/Iso4 isoQTLs for baseline, flu-infected or IFNβ-stimulated cells (bottom 3 panels). Y-axis: −log_10_(P-value) of association. X-axis: genomic location. Points are colored based on their LD to rs1061502. (C) Transcript structure and box-whisker plots (bottom 3 panels) between *IRF7* transcript quantitative traits (y-axis: log2(normalized gene abundance), normalized isoform abundance, or isoform percentage) and rs1061502 genotype (x-axis). (D) Heatmap of genes distally associated (permutation FDR < 0.05) with risoQTL rs1061502. Heatmap colors are row-scaled TPM values (yellow: row max, purple: row min).

The *IRF7* locus harbors an isoQTL within an extended haplotype previously known to be associated with SLE (rs58688157 lead SNP, P < 2.97×10^−11^) (Morris et al. 2016) (**Fig. 4B**). A linked SNP (rs1061502; LD R^2^ = 0.93, D’ = 0.97 to rs58688157) is the most significant association to overall *IRF7* expression in IFNβ-stimulated (P < 2.87×10^−49^) and flu-infected cells (P < 2.21×10^−25^) (**Fig. 4C, Fig. S11**). IsoQTL analysis further revealed that rs1061502^T^ also increased the usage of *IRF7*/Iso4 (flu beta = 5.3%, P < 9.42×10^−52^, IFNβ beta = 5.8%, P < 4.81 ×10^−46^, **Fig. 4C**, panel 3, purple, **Fig. S12**) while decreasing the usage of *IRF7*/Iso3 (flu beta = −3.4%, 1.15×10^−15^, IFNβ beta = −7.0%, 2.04×10^−29^, **Fig. 4C**, panel 3, green, **Fig. S12**). Further, although overall *IRF7* abundances were similar between the two stimulated conditions, *IRF7*/Iso4 (purple) was the dominant isoform in flu-infected cells but not IFNβ-stimulated cells (10.7x fold, P < 10^−306^) (**Fig. 4C**, bottom panel, **Fig. S12**). Rs1061502 was also a distal eQTL (permutation FDR < 0.2) for a cluster of genes including *NMI*, type-1 interferon *IFNA2, IFIT5*, and *C5* only in flu-infected, but not in IFNβ-stimulated cells (**Fig. 4D**). These results replicate and expands our previous findings that rs12805435 (LD R^2^ = 0.95, D’ = 0.98 to rs1061502) is associated in *cis* with *IRF7* expression in IFNβ-stimulated and flu-infected cells, and in *trans* with a cluster of *IRF7*-regulated genes only in flu-infected cells (Lee et al. 2014). The flu-specific *trans* associations could be due to the additive effects of flu-specific induction of *IRF7*/Iso4 independent of IFNβ signaling and induction of overall *IRF7* expression by rs1061502^T^. *IRF7*/Iso4 encodes a 516-amino acid protein product and differs from other dominant isoforms in IFNβ-stimulated cells (*IRF7*/Iso1 and *IRF7*/Iso3) in the 5’ UTR and the coding sequence in the DNA-binding domain. Given the link between a type-1 interferon signature and SLE, this raises an intriguing notion that SNPs affecting a specific *IRF7* isoform could impact viral responses and autoimmune inflammation through similar mechanisms.

### An *ERAP2* risoQTL controls differential transcript usage during influenza infection

The *ERAP2* locus is characterized by two frequent and highly differentiated (40 SNPs in perfect LD) haplotypes observed in every major human population (B: 53% and A: 47%) (**Fig. S13**). The minor haplotype A encodes a 965 amino-acid protein and is associated with Crohn’s disease (Jostins et al. 2012) but not ulcerative colitis (**Fig. 5A**). The major allele (G) of rs2248374, a splice-site variant tagging Haplotype B, creates an alternate 3’ donor splice site inducing the alternative splicing of an extended exon 10 with two premature termination codons (Andres et al. 2010). As a result, transcripts from Haplotype B are degraded by nonsense-mediated decay resulting in one of the most significant eQTLs and isoQTLs in most tissues and cell types (Lappalainen et al. 2013; Lee et al. 2014; Ye et al. 2014; Consortium 2015). Intriguingly, the *ERAP2* locus has been maintained by long-term balancing selection (between 1.4M (Andres et al. 2010) and 5.1M years (Cagliani et al. 2010)) raising the important question: in what environmental conditions do balancing selection act to maintain the seemingly loss-of-function (LOF) Haplotype B and the disease-causing Haplotype A in humans?

**Figure 5.**
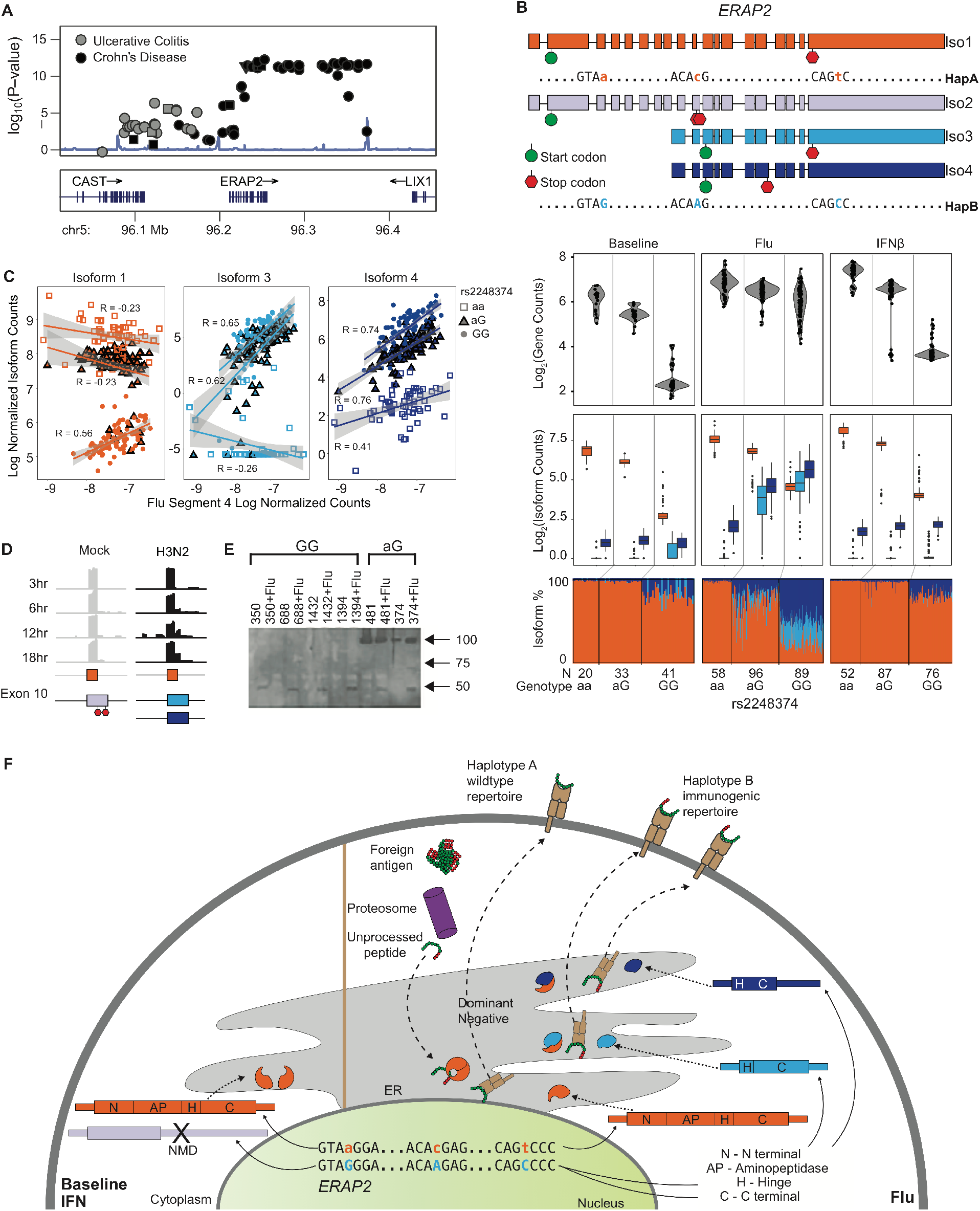
Genetics of *ERAP2* regulation. (**A**) LocusZoom plot of Crohn’s disease and ulcerative colitis associations at the *ERAP2* locus. Y-axis: −log_10_(P-value) of association. X-axis: genomic location. (**B**) Structures of transcripts derived from each haplotype (top panel) and box-whisker plots (3 bottom panels) between *ERAP2* transcript quantitative traits (y-axis: log2(normalized gene abundance), normalized isoform abundance, or isoform percentage) and genotype (x-axis). (**C**) Correlation between *ERAP2*/Iso1 (orange), *ERAP2*/Iso3 (light blue) and *ERAP2*/Iso4 (dark blue) abundance (y-axis) and overall flu transcript abundance (x-axis) segregated by rs2248374 genotype (aa: squares, aG: triangles, GG: circles). (**D**) RNA-seq reads from an independent experiment infecting monocyte derived macrophages with H3N2 provides evidence of time dependent transcription of the short *ERAP2* isoforms. (**E**) Western blot of MoDCs before and after flu-infection from 5 Haplotype B homozygotes and 2 heterozygotes. Full length *ERAP2* protein isoform is expected at 120 kDas. Two flu-specific *ERAP2* protein isoforms are expected at 49 and 29 kDas. (**F**) A schematic of the hypothesized regulation and function of two *ERAP2* haplotypes. N - N terminal domain. AP - amino peptidase domain. H - hinge domain. C - C terminal domain. PTC - premature termination codon.

Given the known role of *ERAP2* in antigen presentation (Saveanu et al. 2005), we examined the genetic control of *ERAP2* transcripts in the human antiviral response. In resting and IFNβ-stimulated cells, we confirmed the known genetic association of rs2248374^G^ allele with lower *ERAP2* expression (**Fig. 5B**). Remarkably, in flu-infected but not IFNβ-stimulated cells, two previously uncharacterized short isoforms (*ERAP2*/Iso3, *ERAP2*/Iso4, **Fig. 5B, Fig. S14**) were transcribed from Haplotype B, resulting in the partial rescue of *ERAP2* expression. The short isoforms differed from the constitutive full-length isoform (*ERAP2*/Iso1 transcribed from Haplotype A) by the initiation of transcription at exon 9 and the alternate splicing of an extended exon 10, and differed from each other by alternative splicing at a secondary splice site at exon 15. The initiation of transcription at exon 9 results in an alternate in-frame translation start site at exon 11 thus rendering the premature termination codon in exon 10 inactive. The influenza-dependent genetic control of *ERAP2* isoform usage is supported by two additional lines of evidence. First, there is significant correlation between overall flu transcript abundance, a proxy for degree of infection, and *ERAP2*/Iso3 and *ERAP2*/Iso4 transcript abundances and usages in heterozygotes and homozygotes for the Haplotype B (**Fig. 5C, Fig. S15**). Second, the transcription of an extended exon 10, a hallmark of flu-specific short isoforms, was observed in monocyte derived macrophages infected by H3N2 over a time course in an independent RNA-seq dataset (fluomics, GEO GSE97672) (**Fig. 5D**).

The predicted protein products of the *ERAP2* short isoforms would be missing the N-terminal (Domain 1) and aminopeptase domains (Domain 2) while maintaining a partial hinge domain (Domain 3) and the full (*ERAP2*/Iso3) or partial (*ERAP2*/Iso4) alpha helical C-terminal domain (Domain 4). This calls into question whether the short *ERAP2* isoforms would function as an RNA or protein product. Western blotting with a C-terminal specific antibody for *ERAP2* isoforms detects at least one short protein isoform (~50 kDa) in flu-infected cells from Haplotype B homozygotes and heterozygotes suggesting the translation of the short influenza-specific *ERAP2* isoforms (**Fig. 5E**).

We present a model of the regulation and function of the *ERAP2* locus consistent with our findings and previous results (**Fig. 5F**). The genetic signals at the *ERAP2* locus suggest at least three perfectly linked variants on Haplotype B affecting *ERAP2* transcription and splicing in response to viral stimulation. Rs2548538, an intronic variant that overlaps chromatin marks from LCLs (Consortium 2012), is a candidate SNP that causes alternate transcript initiation at exon 9. Rs2248374^G^, the known splice site mutation, creates an alternate preferred splice site resulting in alternative splicing of an extended exon 10. Rs2549797^G^, a splice-site mutation that creates a competing alternate splice site, results in ~40% of the transcripts with an extended exon 15. Previous work has shown that full length *ERAP2* is a prototypical aminopeptidase that homodimerizes and heterodimerizes with *ERAP1* (Saveanu et al. 2005) to perform the final peptide trimming step prior to MHC class I loading in the ER. The translation of the flu-specific *ERAP2* isoforms that lack the aminopeptidase domain suggests that it could act as a dominant negative to either *ERAP2* or *ERAP1* to disrupt normal antigen processing creating a more immunogenic MHC peptide repertoire that could confer a fitness advantage in response to virus.

## Discussion

Although maps of genetic variants associated with overall gene abundances have been generated in many tissue types, the genetic control of alternate isoform usage has not been extensively studied. Using *de novo* transcript reconstruction, we found a large number of previously uncharacterized transcripts in human dendritic cells, especially in response to influenza and interferon stimulation, indicating that the current reference human transcriptome is far from complete. We further found genetic variants (isoQTLs) associated with alternate isoform usage are widespread, > 40% of which are not associated with the overall abundance of the corresponding gene, indicative of independent genetic control of gene regulation at these loci. The enrichment of isoQTLs for known splice sites and autoimmune and neurodegenerative disease loci suggest a highly clinically relevant set of candidate variants that induce targetable changes in protein sequence.

IsoQTLs, like eQTLs, can affect gene expression at other loci in the genome suggesting important downstream effects on gene regulation. The most striking example is at the *IRF7* locus where a splice-site SNP affects *IRF7* splicing in response to influenza and interferon, but only affects the expression of downstream genes in response to flu. This suggests that both genetic effects on isoform usage and stimulation dependent regulation of *IRF7* expression are necessary for the observed *trans* effects. Although C-terminal splice forms of *IRF7* have been shown to differentially transactivate type-1 interferons and chemokines (Lin et al. 2000), IRF7/Iso4 is not known to have specific antiviral properties *in vivo* even though its ectopic expression is known to activate IFNAs in fibroblasts (Au et al. 1998). The association of the variant with SLE indicates a possible role for viral exposure to prime the immune system of individuals carrying the risk allele toward autoimmunity.

Different genetic variants in a locus could affect multiple facets of gene regulation in response to stimulation in establishing transcriptional diversity. This was clearly demonstrated at the *ERAP2* locus where multiple variants lead to differential expression and splicing of short isoforms in response to influenza but not interferon. The lack of expression in response to IFNβ suggests the transcription of novel *ERAP2* isoforms is likely initiated by viral sensing pathways upstream of type 1 interferon signaling. Furthermore, balancing selection at *ERAP2* suggests that transcripts derived from both haplotypes could confer fitness advantages, likely in different environments. We provide evidence of the expression of short *ERAP2* isoforms encoded by Haplotype B in response to influenza suggesting viral infection as a possible selective agent. One can speculate that the long *ERAP2* isoform encoded by Haplotype A could be selected under different environmental conditions that favors an overactive autoinflammatory response or a primed T cell response. Altogether, the genetic analysis of isoforms under physiologically relevant conditions can help reveal new gene regulatory mechanisms by which alleles associated with disease and under natural selection function in response to environment.

## Methods

### Study subjects

Donors were recruited from the Boston community and gave written informed consent for the studies. Individuals were excluded if they had a history of inflammatory disease, autoimmune disease, chronic metabolic disorders or chronic infectious disorders. Donors were between 18 and 56 years of age (avg. 29.9).

### Reagents

Influenza A (PR8 ΔNS1) was prepared as described (Shapira et al. 2009). Recombinant human IFNβ was obtained from PBL Assay Science (Piscataway, NJ). Antibodies used were anti-IRF1 (sc-497x; Santa Cruz Biotechnology; Dallas, TX), anti-STAT2 (sc-476x; Santa Cruz Biotechnology) and anti-IRF9 (sc-10793x; Santa Cruz Biotechnology).

### Preparation and stimulation of primary human monocyte-derived dendritic cells

As previously described(Lee et al. 2014), 35-50 mL of peripheral blood from fasting subjects was collected between 7:30-8:30 am. The blood was drawn into sodium heparin tubes and peripheral blood mononuclear cells (PBMCs) were isolated by Ficoll-Paque (GE Healthcare Life Sciences; Uppsala, Sweden) centrifugation. PBMCs were frozen in liquid N_2_ in 90% FBS (Sigma-Aldrich; St. Louis, MO) and 10% DMSO (Sigma-Aldrich). Monocytes were isolated from PBMCs by negative selection using the Dynabeads Untouched Human Monocytes kit (Life Technologies; Carlsbad, CA) modified to increase throughput and optimize recovery and purity of CD14+CD16^lo^ monocytes: the FcR Blocking Reagent was replaced with Miltenyi FcR Blocking Reagent (Miltenyi; Bergisch Gladbach, Germany); per mL of Antibody Mix, an additional 333 ug biotinylated anti-CD16 (3G8), 167 ug biotinylated anti-CD3 (SK7) and 167 ug biotinylated anti-CD19 (HIB19) antibodies (Biolegend; San Diego, CA) were added; the antibody labeling was modified to be performed in 96-well plates; and Miltenyi MS Columns or Multi-96 Columns (Miltenyi) were used to separate magnetically-labeled cells from unlabeled cells in an OctoMACS Separator or MultiMACS M96 Separator (Miltenyi) respectively. The number of PBMCs and monocytes was estimated using CellTiter-Glo Luminescent Cell Viability Assay (Promega; Madison, WI). A subset of the isolated monocytes was stained with PE-labeled anti-CD14 (M5E2; BD Biosciences; Franklin Lakes, NJ) and FITC-labeled anti-CD16 (3G8; Biolegend), and subjected to flow cytometry analysis using an Accuri C6 Flow Cytometer (BD Biosciences). A median of 94% CD14+ cells and 99% CD16^lo^ cells was obtained. The remaining monocytes were cultured for seven days in RPMI (Life Technologies) supplemented with 10% FBS, 100 ng/mL GM-CSF (R&D Systems; Minneapolis, MN) and 40 ng/mL IL-4 (R&D Systems) to differentiate the monocytes into monocyte-derived dendritic cells (MoDCs). 4×10^4^ MoDCs were seeded in each well of a 96-well plate, and stimulated with influenza virus for 10 h, 100 U/mL IFNβ for 6.5 h or left unstimulated. Cells were then lysed in RLT buffer (Qiagen; Hilden, Germany) supplemented with 1% β-mercaptoethanol (Sigma-Aldrich).

### RNA isolation and sequencing

RNA from all samples was extracted using the RNeasy 96 kit (Qiagen, cat. # 74182), according to the manufacturer’s protocols. 576 total samples were sequenced (99 baseline, 250 influenza infected and 227 interferon stimulated). 552 pass filter samples (94 baseline, 243 influenza and 215 interferon) were sequenced to an average depth of 38M 76bp paired end reads using the Illumina TruSeq kit. Reads were aligned to hg19 genome with Tophat v1.4.1 (--mate-inner-dist 300 --mate-std-dev 500)(Trapnell et al. 2009) with 86% mapping to transcriptome and 97% mapping to the genome (**Table S1**).

### Adjusting for expression heterogeneity

We empirically determined the number of principal components to adjust for each stimulation condition and either overall gene abundance or isoform percentage. Because of the smaller number of individuals in the baseline study, the number of principal components adjusted is fewer (**Fig. S2**). Because the isoform percentage implicitly adjusts for confounders that affect overall gene abundance and isoform abundance levels (i.e. other eQTLs), the number of adjusted PCs is also fewer (**Fig. S3**).

### Transcriptome reconstruction

After aligning reads to the genome, we reconstructed transcriptomes for each sample individually using StringTie (Pertea et al. 2015) using default parameters and quantified the abundances of annotated transcripts using kallisto (Bray et al. 2016). For genes expressed at > 5 transcripts per million (TPM) in any sample, we removed isoforms expressed at < 5 TPM across all samples. In order to preserve isoforms that may be uniquely expressed in a single condition (e.g. baseline, flu, IFN), transcriptomes within the same conditions were first merged before transcriptomes across all three conditions were merged, using cuffcompare (Trapnell et al. 2012). As a final step, cuffmerge (Trapnell et al. 2012) (--overhang-tolerance 0) was used to remove redundant isoforms.

### Transcriptome comparison

We compared our *de novo* reconstructed isoforms with the compendium of annotated isoforms from Gencode (v27) (Harrow et al. 2006), UCSC (Casper et al. 2018) (hg19), and RefSeq (O’Leary et al. 2016) (hg19). For each reconstructed isoform, we compared the position of each splice site as well as the 5’ and 3’ position of each splice junction to all annotated isoforms, and report comparison statistics for the most similar annotated isoform.

To quantify the coverage of spliced reads across splice junctions, we fed each tophat alignment to leafcutter (Li et al. 2018), which uses the CIGAR strings in alignment .bam files to count the number of high-quality aligning reads at each splice junction.

To detect novel TSSs, we overlapped the first 100 bp of our reconstructed isoforms with TSSs of annotated isoforms. We further characterized the novel TSSs detected in our dataset by comparing to CAGE reads from MoDCs from the Fantom5 database (Noguchi et al. 2017) (FF:11227-116C3, F:11308-117C3, FF:11384-118B7). Using our *de novo* isoforms, we created a specialized transcriptome consisting of only the first 500 sequences of each transcript (stranded, and after splicing). We then aligned CAGE reads to this specialized transcriptome using bowtie v0.12.7 with default parameters (Langmead 2010), and quantified the presence of each TSS by counting the number of mapped reads.

### Transcriptome quantification

Differential expression testing was carried out with sleuth (Pimentel et al. 2017) using 100 bootstraps per sample. Gene-level quantification was estimates by summing isoform counts and differential expression testing was carried out with DESeq2 (Anders and Huber 2010).

### Beta regression

Differential isoform ratio testing was carried out in R using beta regression package betareg (Cribari-Neto and Zeileis 2010) and P-values were calculated using likelihood ratio test and adjusted with a false discovery rate adjustment.

### Gene ontology (GO) enrichment analysis

GO enrichment analysis was carried out using Gorilla (Eden et al. 2009) and tested against a background of only the set of genes that were expressed in monocyte derived dendritic cells and which we recovered during the transcriptome reconstruction (see above).

### DNA extraction and genotyping

As previously described (Lee et al. 2014), genomic DNA was extracted from 5 mL whole blood (DNeasy Blood & Tissue Kit; Qiagen), and quantified by Nanodrop. Each subject was genotyped using the Illumina Infinium Human OmniExpress Exome BeadChips, which includes genome-wide genotype data as well as genotypes for rare variants from 12,000 exomes as well as common coding variants from the whole genome. In total, 951,117 SNPs were genotyped, of which 704,808 SNPs are common variants (Minor Allele Frequency [MAF] > 0.01) and 246,229 are part of the exomes. The genotype success rate was greater than or equal to 97%. We applied rigorous subject and SNP quality control (QC) that includes (1) gender misidentification, (2) subject relatedness, (3) Hardy-Weinberg Equilibrium testing, (4) use concordance to infer SNP quality, (5) genotype call rate, (6) heterozygosity outlier, (7) subject mismatches. In the European population, we excluded 1,987 SNPs with a call rate < 95%, 459 SNPs with Hardy-Weinberg equilibrium P-value < 10^−6^, 234 SNPs with a MisHap P-value < 10^−9^, and 63,781 SNPs with MAF < 1% from (a total of 66,461 SNPs excluded). In the African-American population, we excluded 2,161 SNPs with a call rate < 95%, 298 SNPs with Hardy-Weinberg equilibrium P-value < 10^−6^, 50 SNPs with a MisHap P-value < 10^−9^, and 17,927 SNPs with MAF < 1% from (a total of 20,436 SNPs excluded). In the East Asian population, we excluded 1,831 SNPs with a call rate < 95%, 213 SNPs with Hardy-Weinberg equilibrium P-value < 10^−6^, 47 SNPs with a MisHap P-value < 10^−9^, and 84,973 SNPs with MAF < 1% from (a total of 87,064 SNPs excluded). After QC, 52 subjects across all three populations and approximately 18,000 – 88,000 SNPs in each population were filtered out from our analysis.

Population stratification: Underlying genetic stratification in the population was assessed by multidimensional scaling using data from the International HapMap Project (International HapMap 2003) (CEU, YRI and CHB samples) combined with IBS cluster analysis using the Eigenstrat 3.0 software(Price et al. 2006).

The quality control of the genotyping data were performed using PLINK (Purcell et al. 2007).

### Genotype imputation

To accurately evaluate the evidence of association signal at variants that are not directly genotyped, we used the BEAGLE (Browning and Browning 2016) software (version: 3.3.2) to imputed the post-QC genotyped markers using reference Haplotype panels from the 1000 Genomes Project (Siva 2008) (The 1000 Genomes Project Consortium Phase I Integrated Release Version 3), which contain a total of 37.9 Million SNPs in 1,092 individuals with ancestry from West Africa, East Asia, and Europe. For subjects of European and East Asian ancestry, we used haplotypes from Utah residents (CEPH) with Northern and Western European ancestry (CEU), and combined panels from Han Chinese in Beijing (CHB) and Japanese in Tokyo (JPT), respectively. For imputing African American subjects, we used a combined haplotype reference panel consisting of CEU and Yoruba in Ibadan, Nigeria (YRI). For the admixed African American population, using two reference panels substantially improves imputation performance. After genotype imputation, we filtered out poorly imputed (BEAGLE *r*^2^ < 0.1) and low MAF SNPs (MAF < 0.01), which resulted in 7.7M, 6.6M, 12.7M common variants in European, East Asian and African American, respectively. This set of genotyped and imputed markers was used for all the subsequent association analysis.

### eQTL and isoQTL mapping

QTL mapping was performed using the Matrix eQTL (Shabalin 2012) package using empirically determined number of principal components (PCs) as covariates for each analysis. For *cis* QTLs, isoform usage or overall gene abundance were regressed against all genetics variants with a MAF > 5% in a 1MB (+/-500kb) window and the most significant association is kept. For *cis* QTLs, 0 – 44 PCs (*cis* eQTLs) and 0 – 12 PCs (*cis* isoQTLs) in increments of 2 were tested and the number of PCs were chosen to maximize the number of *cis* e/isoQTLs detected (**Fig. S2** and **S3**). Experiment wide empirical P-values were calculated by comparing the nominal P-values with null P-values determined by permuting each isoform/gene 1,000 times (Churchill and Doerge 1994). The permutation p values are not pooled to calculate the empirical p values (i.e. the minimum possible P-value is 0.001). False discovery rates were calculated using the qvalue package (JDSwcfAJ et al. 2015) as previously described (Storey and Tibshirani 2003).

### reQTL and risoQTL mapping

QTL mapping was performed using the Matrix eQTL (Shabalin 2012) package using empirically determined number of principal components (PCs) as covariates for each analysis. For *cis* QTLs, change in isoform usage or overall gene abundance were regressed against all genetics variants with a MAF > 5% in a 1MB (+/-500kb) window and the most significant association is kept. For *cis* QTLs, 0 – 44 PCs (*cis* reQTLs) and 0 – 12 PCs (*cis* risoQTLs) in increments of 2 were tested and the number of PCs were chosen to maximize the number of *cis* re/risoQTLs detected (**Fig. S2** and **S3**). Experiment wide empirical P-values were calculated by comparing the nominal P-values with null P-values determined by permuting each isoform/gene 1,000 times(Churchill and Doerge 1994). Because of the lower number of true positives, the permutation p values were pooled to calculate the empirical p values in order to obtain robust estimates of q values. False discovery rates were calculated using the qvaluepackage (JDSwcfAJ et al. 2015) as previously described (Storey and Tibshirani 2003).

### QTL annotation

QTLs were annotated using Variant Effect Predictor and Ensembl release 79(McLaren et al. 2016). Exonic and intronic locations of QTLs were determined using UCSC’s canonical transcripts (table *knownCanonical)* as a reference (Karolchik et al. 2004). Enrichments were calculated against background set of SNPs that were matched in allele frequency (binned by 4%) and distance to nearest transcription start site (binned by 10kb).

### Overlap with GWAS associations

The GREGOR suite (Schmidt et al. 2015) was used for calculating the enrichment of eQTLs and isoQTLs containing a GWAS loci across baseline, flu, and IFNβ stimulations. GWAS associations for disease with FDR < 0.1 are reported.

### Partitioned heritability analysis

We used LD score regression (Finucane et al. 2015) to calculate the proportion of SNP heritability explained by eQTL/isoQTLs. We obtained summary statistics from 28 human traits/diseases from https://data.broadinstitute.org/alkesgroup/sumstats_formatted/ and ran ldsc with default parameters.

### Estimating flu transcript abundance

Flu transcript abundance was estimated by running RSEM (Li and Dewey 2011) on a custom reference of the influenza PR8 genome.

### ERAP2 western blot

Protein extracts were fractionated by SDS-PAGE (4-12% Bis-Tris gel, Thermo scientific, NP0335BOX) and transferred to PVDF membrane (BioRad, cat. #162-0177). After blocking with 2% BSA in TBST (Tris buffered saline containing 0.1% tween-20) for 1 hour, membranes were incubated with primary antibody (either ERAP2, R&D Systems, cat# AF3830, 1:3000) or b-actin (Abcam, cat. #ab6276, 1:15,000) overnight at 4C. Membranes were then washed and incubated with a 1:5000 dilution of HRP conjugated secondary antibody (either donkey anti-goat from Santa Cruz Biotech cat. #sc2020, or with goat anti-mouse from Jackson immune Research cat. #115-035-146) for 1 hour. Membranes were washed and developed with ECL system (VWR, cat. #89168-782) according to the manufacturer’s protocol.

## Data Access

Processed RNA-Sequencing data is available from GEO under accession GSE92904. Raw fastq data is available from dbGAP under accession phs000815.v1.p1.

